# Automated Summarisation of SDOCT Volumes using Deep Learning: Transfer Learning vs *de novo* Trained Networks

**DOI:** 10.1101/424507

**Authors:** Bhavna J. Antony, Stefan Maetschke, Rahil Garnavi

## Abstract

Spectral-domain optical coherence tomography (SDOCT) is a non-invasive imaging modality that generates high-resolution volumetric images. This modality finds widespread usage in ophthalmology for the diagnosis and management of various ocular conditions. The volumes generated can contain 200 or more B-scans. Manual inspection of such large quantity of scans is time consuming and error prone in most clinical settings. Here, we present a method for the generation of visual summaries of SDOCT volumes, wherein a small set of B-scans that highlight the most clinically relevant features in a volume are extracted. The method was trained and evaluated on data acquired from age-related macular degeneration patients, and “relevance” was defined as the presence of visibly discernible structural abnormalities. The summarisation system consists of a detection module, where relevant B-scans are extracted from the volume, and a set of rules that determines which B-scans are included in the visual summary. Two deep learning approaches are presented and compared for the classification of B-scans - transfer learning and *de novo* learning. Both approaches performed comparably with AUCs of 0.97 and 0.96, respectively, obtained on an independent test set. The *de novo* network, however, was 98% smaller than the transfer learning approach, and had a run-time that was also significantly shorter.

## 1 Introduction

The detection of key frames is a common approach employed in video analysis, particularly for the summarisation of video sequences. The techniques typically rely on the detection of explicit features of interest such as motion [1, 2] as well as other features such as edge information [3] and self-similarity [4]. Condensing videos via shot boundary detection has also been applied to the summarisation of video sequences [5].

In medical imaging, the detection of keyframes is more commonly found in the analysis of angiogram video sequences, but is less common in other radiographic modalities. Gibson *et al*. [6] described an approach for the compression of angiogram videos by detecting diagnostically relevant frames in videos and ensuring they were preserved. Syeda-Mahmood *et al*. [7] presented an approach for the detection of key frames in angiogram video analysis by detecting the vessels and selecting frames in which their visibility was best. However, both of these approaches relied on the explicit detection of features of interest in the images in order to identify them as “key”.

In ophthalmology, spectral-domain optical coherence tomography (SDOCT) [8] has begun to find widespread use for the diagnosis and management of various ocular conditions. This non-invasive imaging modality relies on laser interferometry to generate high-resolution images of the retina, which allows for the visualisation and quantification of structures in 3-D. These volumetric images are comprised of B-scans, which numbers can range from as few as five to two hundred or more. Summarisation of these volumes in current scanning systems is usually limited to a report that indicates the thicknesses of retinal layers. While such a report shows large pathologies such as choroidal neovascularizations (CNV), smaller abnormal indicators such as drusen, epiretinal membranes and microcytic macular edema would not be visible. Thus, visual summaries could complement the existing approach, by highlighting the pathological conditions that are currently not quantified. Previously, Chakravarthy *et. al* [9] described an approach for the detection of B-scans that show choroidal neovascularization (CNV). The method relies on the detection of the retina, followed by a machine-learning approach for the detection of possible fluid patches in the images. While this approach does in fact extract the specific B-scans, the method is limited to CNVs associated with wet-AMD.

Here, we present a deep learning approach for the automated summarisation of SDOCT volumes. Similar to previous summarisation techniques our proposed system begins with the detection of “key” B-scans. The system was trained and tested on SDOCT volumes acquired from patients that presented with age-related macular degeneration (AMD), and “relevance” was defined on the basis of the presence of visibly discernible structural abnormalities. Using a deep learning approach for this task allows it to be posed as a recognition task, and thus, does not require the explicit extraction of features (such as CNV) in order to characterise the B-scan. We employed and compared two deep learning techniques for keyframe extraction, where one is a transfer learning technique based on a pretrained network, while the second is a *de novo* trained custom convolutional neural network (CNN) that is significantly smaller. Transfer learning is a commonly used technique that allows for the repurposing of pretrained networks in applications where data might be scarce (as is commonly the case in medical imaging). Once the relevant B-scans had been identified, a set of rules were applied to generate the visual summary.

The paper is organised as follows: Section 2 details the data used in this experiment; Section!3 describes the two deep learning networks as well as the summarisation rules. The evaluation and comparison of the two networks is presented in Section!4, and a final discussion of the results can be found in Section 5.

## 2 Data

The data used in the experiments were SDOCT images acquired as part of the AREDS2 Ancilliary Study. As detailed in [10], the dataset was registered at ClinicalTrials.gov (Identifier: NCT00734487) and approved by the institutional review boards at 4 A2A SDOCT clinics. With adherence to the tenets of the Declaration of Helsinki, informed consent was obtained from all subjects.

The study cohort consisted of 115 healthy individuals and 269 patients with age-related macular degeneration (AMD). The images were acquired on a Bioptigen SDOCT scanner (Leica Microsystems Inc., Illinois) from an approximately 6.7×6.7mm area centred on the fovea. Each volumes consisted of 100 B-scans, each containing 1000 A-scans and 512 pixels per A-scan (see Fig. 1(a)). Further details of the study are provided in [10].

**Figure 1.**
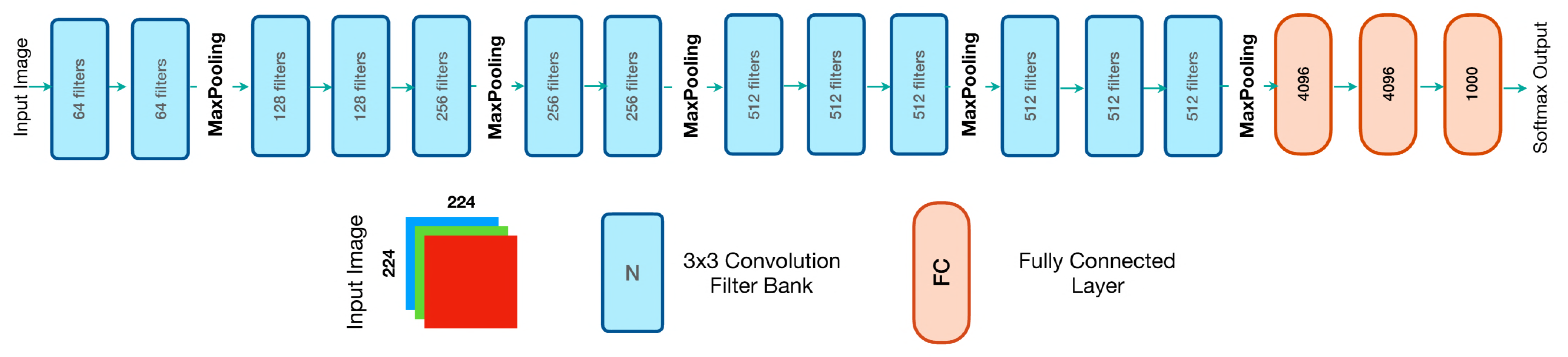
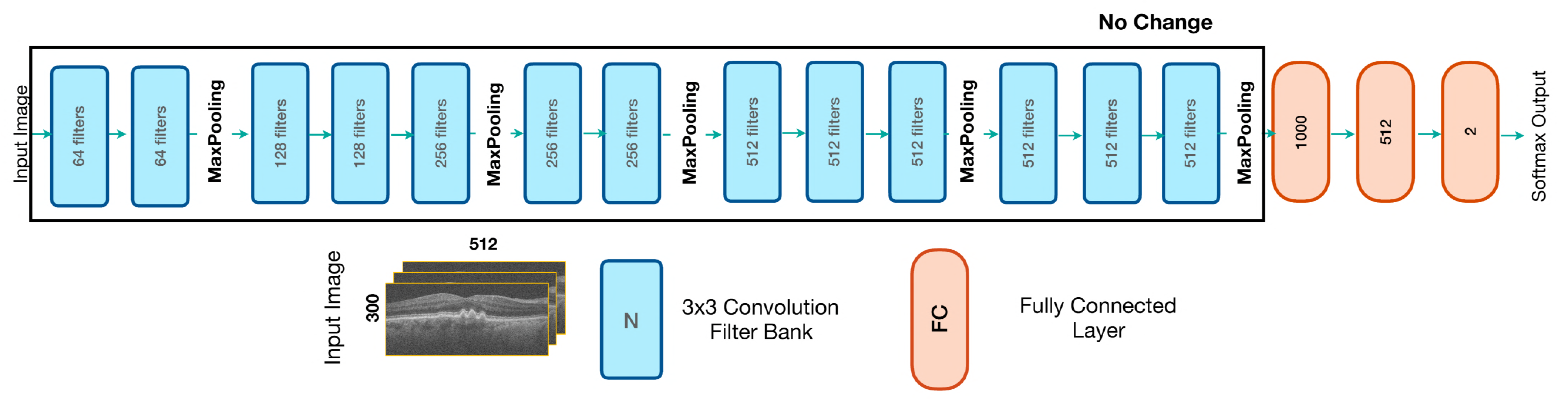
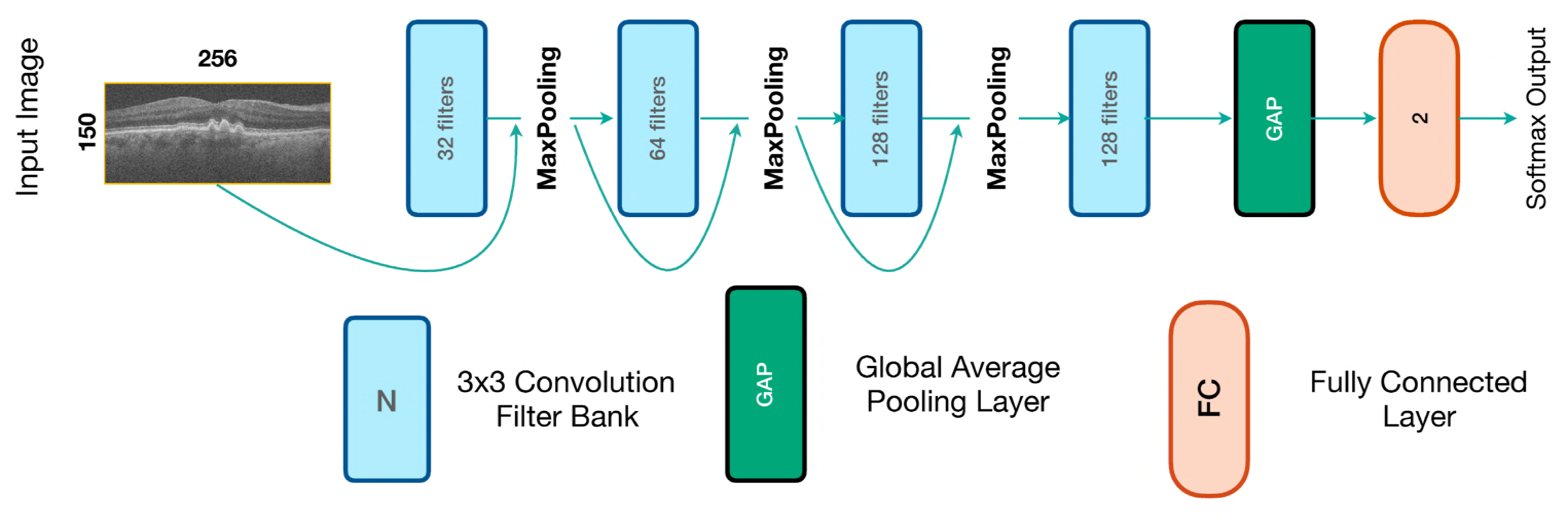
(a) The dimensions of an SDOCT volume depicted on the en-face image (left) and the central B-scan of the volume (right). Examples of poor quality scans with (b) poor contrast, (c) large shadows, and (d) incorrect mirror position.

### 2.1 Data Annotation

The volumes were manually annotated and labeled as being healthy, relevant (containing visibly discernible structural change) or low-quality. For this, each B-scan was visualised and labeled relevant if any visual structural change was observed. Thus, retinal layer thinning (which is difficult to identify visually) was not considered a key feature. However, B-scans with even minor disruptions like small drusen, reticular pseudodrusen and epiretinal membranes were all labeled as relevant “key” B-scans.

The presence of large shadows, poor contrast and other artefacts (such as vinetting, mirror location errors) were flagged as poor quality B-scans (see Fig. 1(b)-(d)).

## 3 Methods

Deep learning [11] has been successfully employed for a number of applications in computer vision such as image recognition [12, 13] and semantic segmentation [14, 15]. This technique has also found application in medical imaging [16] for recognition [17], segmentation [18–20] as well as image registration [21–23]. While larger architectures have shown to perform better than shallower networks, their training also requires larger datasets.

Transfer learning is a technique that re-purposes existing, trained models for new tasks by retraining only small parts of the network. As most weights of the network are left unchanged, this reduces the amount of training data required. Transfer learning lends itself to medical imaging quite well, as large datasets are difficult to acquire in the medical domain. Thus, this was the first technique we employed for the detection of relevant B-scans (detailed in Section 3.1). We utilised the 16-layer VGG network [13] that was initially trained for the ImageNet Challenge - a classification problem consisting of 1000 classes of natural scene images [24].

Transfer learning, however, is not without problems. The pre-trained networks were designed for object recognition in natural scene images, and require the input to be a 3-channel RGB image. SDOCT images are however, grayscale. Thus, a B-scan either has to be replicated three times to meet the required input dimensions, or a section of three slices (and only three) has to be used as network input. Designing and training a network *de novo* allows to circumvent this requirement and potentially could also result in a smaller or more accurate network. For comparison we therefore also designed a significantly smaller convolutional neural network (CNN) that was trained *de novo* (detailed in Section 3.2).

The networks were developed in Keras [25] with TensorFlow [26] as the backend and nuts-flow/ml [27] for the data pre-processing.

### 3.1 Transfer Learning Approach

The structure of the 16-layer VGG network [13] is as shown in Fig. 2(a). It consists of 3×3 convolutional filters with a stride of 1; padded to preserve spatial resolution. All layers utilised rectified linear unit [12] (ReLU) activation. The original network for an input RGB (3-channel) image of 224−224 pixels in size contained 138 million parameters. See [13] regarding training of the original network.

**Figure 2.**
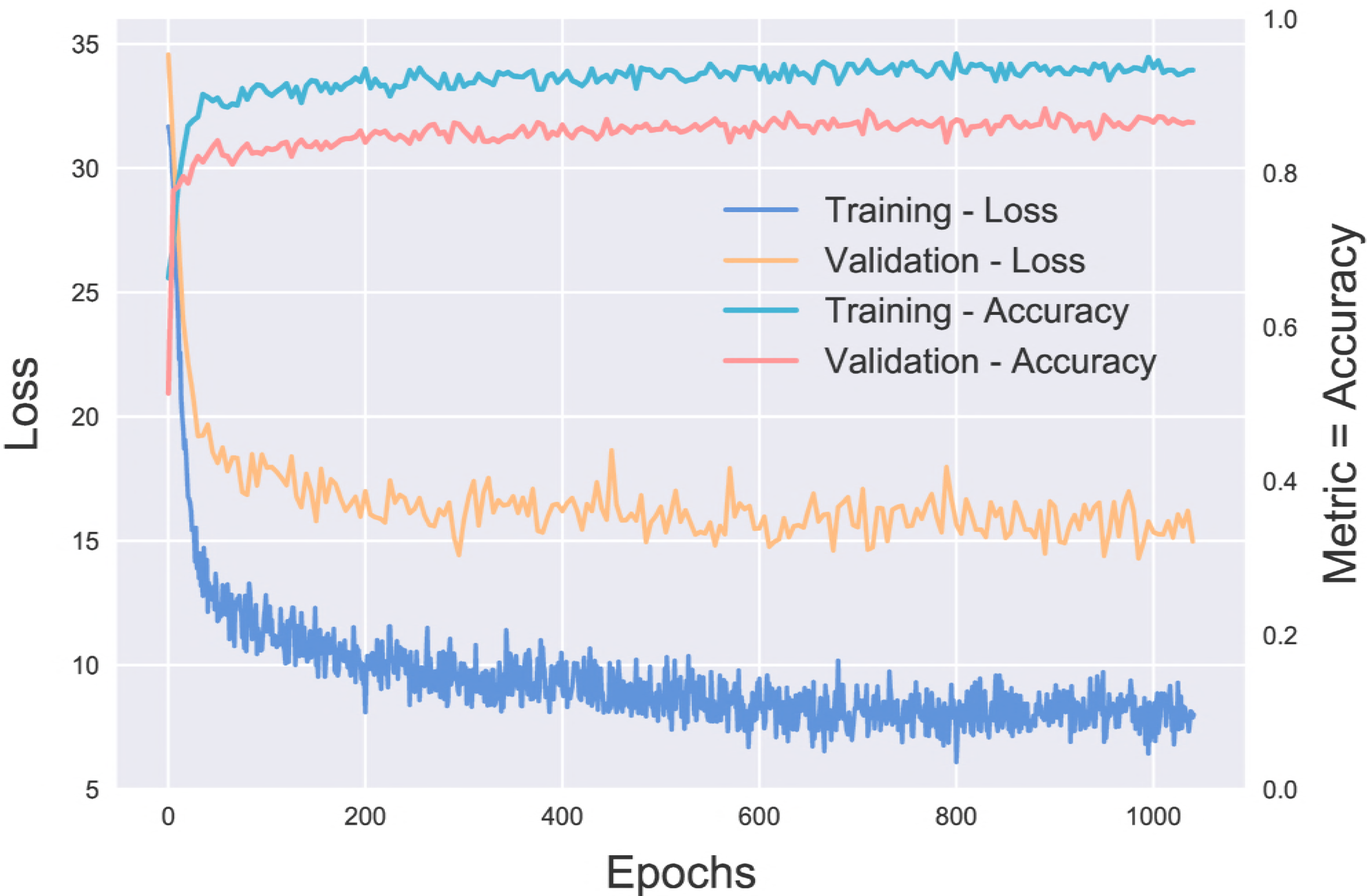
The structure of the (a) original VGG-16 network [13], and (b) the modified transfer network for keyframe detection. Note that the layers in the first five banks of CNNs have not been changed from the original network, and that only the last three fully-connected layers were re-trained.

Since the classification task at hand is a 2-class problem, the model was changed to reflect this (see Fig. 2(b)). Furthermore, the two fully connected layers prior to the final layer were reduced in size from 4096 to 1024 and 512, respectively. Removal of the two original fully connected layers allowed for the input size of the network to be changed to 300×512×3 pixels. For an input image of this size, the total size of the network is 18.6 million parameters. The five blocks of convolutional filters were used as feature extractors and were not re-trained or fine-tuned, resulting in a network with 3.8 million trainable parameters.

Network weights were opimized by Adam [28], with parameters set to recommended values (learning rate set to 1^−6^, *β*_1_ = 0.9, *β*_2_ = 0.999, and *∈* = 1^−8^). The loss function was the balanced cross-entropy loss function:

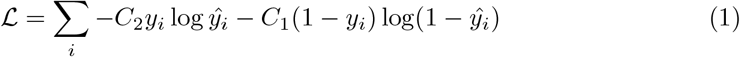

where, *y_i_* is the true label and *ŷ_i_* is the predicated label of the *i*-th sample, and *C*_1_ and *C*_2_ are the number of samples of the first and second class in the batch, respectively. This loss function, being normalised by the number of samples in each class helps with class imbalances. Training was stopped when the validation loss did not decrease by more than 0.1 or after 150 epochs.

#### Data preprocessing

The individual B-scans in each volume are 512×1024 pixels in size. The retina however, does not encompass the entire B-scan, with a large portion of the image showing the vitreous, choroid and scleral tissue. Thus, detecting and extracting the image region that contains the retina reduces the image size. Therefore, the image was filtered with a gaussian derivative filter (first order, *σ*=6.0). This generated a high response at the internal limiting membrane (ILM), and the ellipsoid zone of the photoreceptors as shown in Fig. 3(a)-(b). This response was then thresholded (using a threshold obtained by Otsu’s method [29]), and the largest connected components were detected. Since the largest two components belong to the retinal surfaces, their locations defined the bounding boxes (at least 300 pixels in height) around the retina, and B-scan were cropped to this size. The cropped B-scan was finally resized to 300×512, and replicated three times to match the requirements of the VGG-16 network, which expects the input to be a 3-channel image. The training data was augmented through random rotations (±5°), translations (±10 pixels), contrast scaling (0.3, 1.7) and flipping along the horizontal axis.

**Figure 3.**
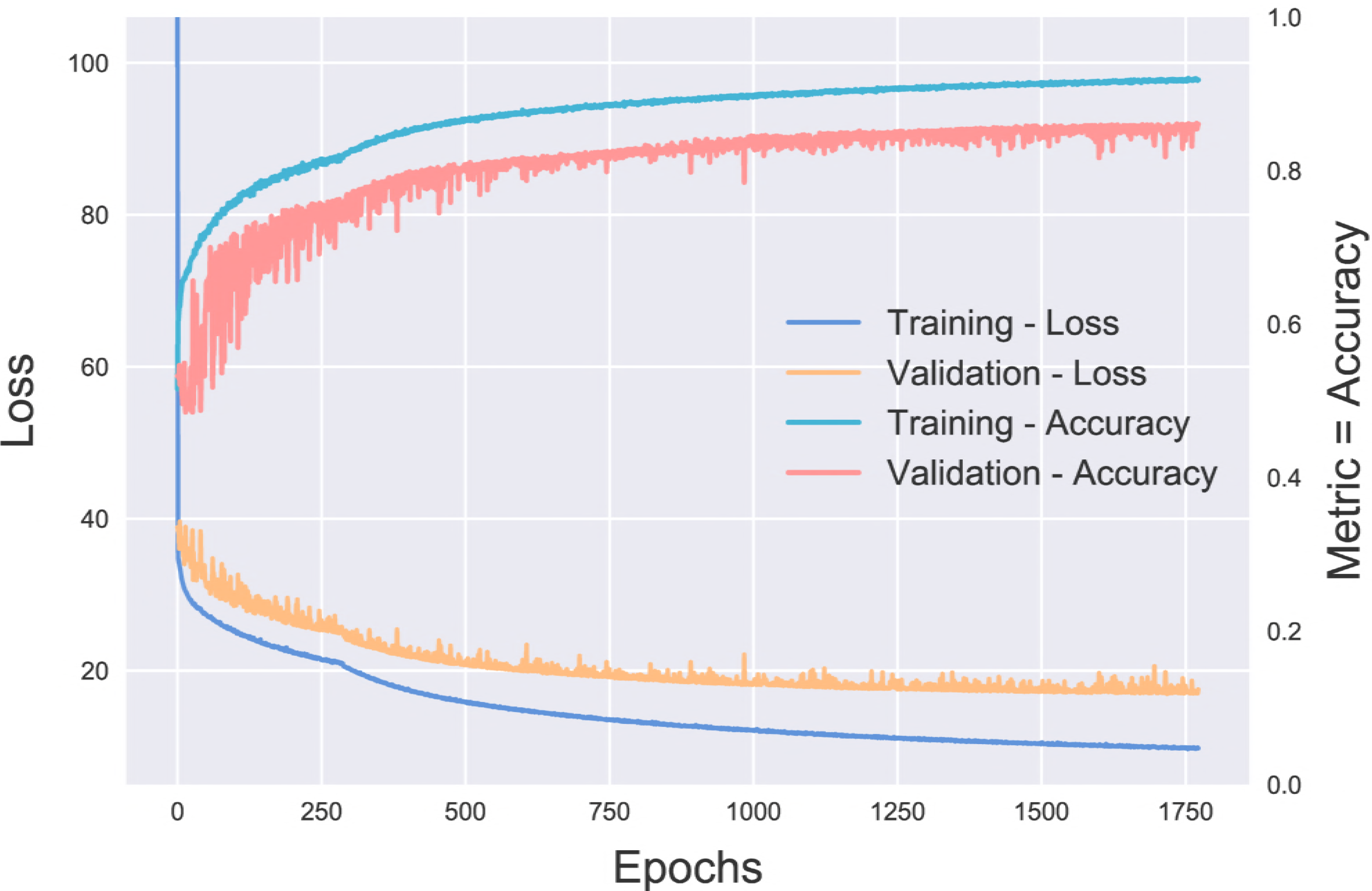
Outline of the data preprocessing steps showing (a) the original slice, (b) the filtered image, (c) connected components of the thresholded image, and (d) the final cropped image indicated by the red box.

### 3.2 De Novo Network

The *de novo* CNN consisted of 4-layers with 32, 64, 128, and 128 filters (3×3 in size). Skip connections [30] were introduced between the layers, with the outputs from the previous layers being concatenated prior to pooling. ReLU activation, and pooling (maximum in a 2×2 window) followed each CNN layer (see Fig. 4). A global average pooling (GAP) layer was added to enable the generation of class activation maps (CAM) [31]. Finally, a fully connected layer (size = 2) with a softmax output provided the class probability for the input B-scan. The resulting network contained only 391300 trainable parameters, and is 2% the size of the transfer learning network. As before, training was performed by Adam with the balanced cross-entropy loss function (see Eq. 1).

**Figure 4.**
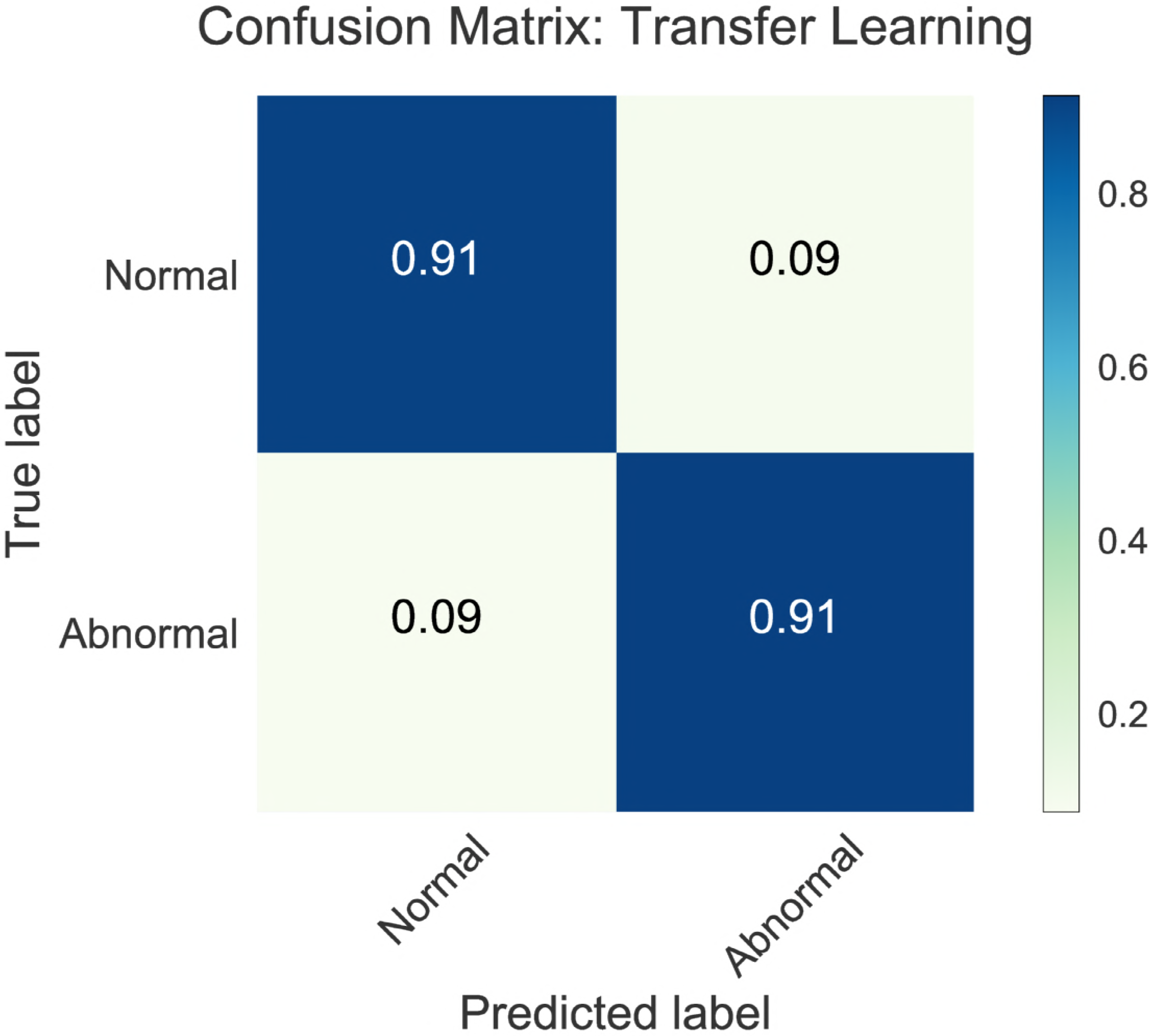
Architecture of the *de novo* network for keyframe detection.

#### Data Preprocessing

The individual B-scans were processed as described for the transfer-learning approach, beginning with the detection of the retina followed by image cropping at the bounding box. The resulting images were down-sampled by a factor of 2 (final size 150×256 pixels) and directly inputted into the network as 1-channel gray-scale images. Training data was augmented as before, employing random rotations (± 5°), translations (± 10 pixels), contrast scaling (0.3, 1.7) and flipping along the horizontal axis.

### 3.3 Experimental Setup

This annotated dataset was then divided into training, validation and testing sets containing 75%, 10% and 15% of the SDOCT volumes, respectively. The allocation of an entire volume to a set ensured that B-scans from a single volume were not distributed across the sets. The final numbers of B-scans - healthy and relevant - in each set are as shown in Table 1.

**Table 1.**
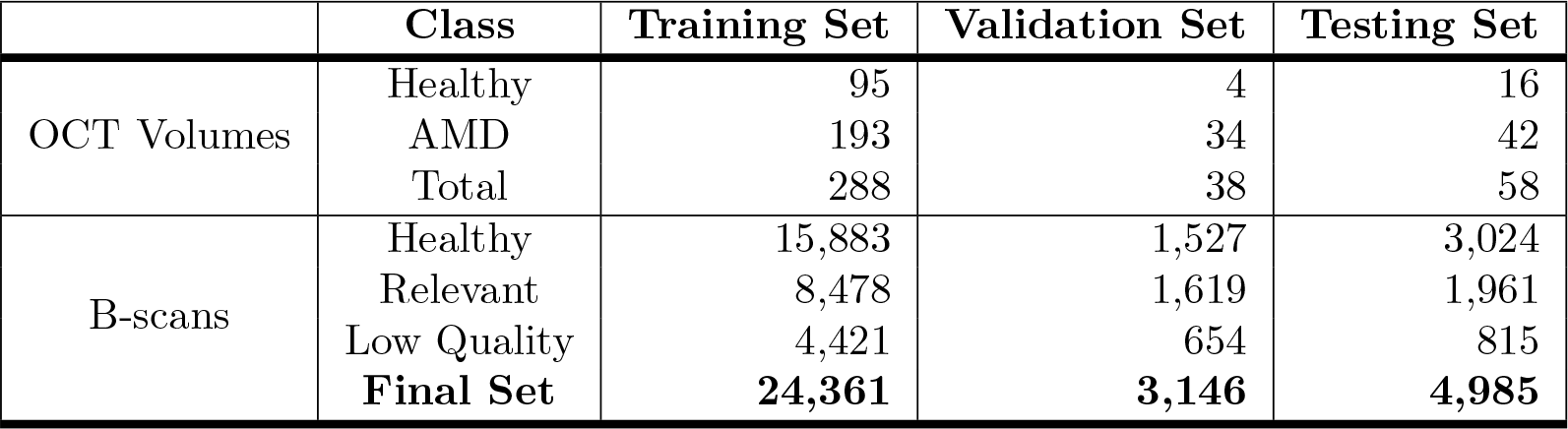
Sizes of training, validation and testing sets.

|  | Class | Training Set | Validation Set | Testing Set |
| --- | --- | --- | --- | --- |
| OCT Volumes | Healthy | 95 | 4 | 16 |
|  | AMD | 193 | 34 | 42 |
|  | Total | 288 | 38 | 58 |
| B-scans | Healthy | 15,883 | 1,527 | 3,024 |
|  | Relevant | 8,478 | 1,619 | 1,961 |
|  | Low Quality | 4,421 | 654 | 815 |
|  | **Final Set** | **24,361** | **3,146** | **4,985** |

The performance of the two networks was evaluated using the area under the curve (AUC). The false positive and false negative rates were also computed and compared for the two networks.

### 3.4 Summarisation Rules

Once the relevant B-scans have been extracted, a set of rules is imposed to select the key-frames (see Algorithm 1). If no relevant B-scans were identified by the deep learning framework, then three slices (two peripheral and one central slice) are returned by the system. Otherwise, the function *RegionDetector()* analyses the set of relevant B-scans *F_i_*, *i* = 1, 2, *… N*, and groups them into regions. If two of the identified B-scans are only separated by a small distance (preset threshold *T*), they are considered to part of the same region. This not only compensates for mislabeled B-scans (not correctly identified as relevant for the summary), but also aggregates small regions that show disease-induced change. For instance, drusen may be present in a small number of B-scans near the fovea, but the individual slices may be separated by a few slices that show no pathology. In such a situation, it is reasonable to aggregate them into a larger region. A flexible threshold *T*, controls the aggregation during run time.

Next, each region is represented by the first, median and last scan of each region. Thus, if *M* regions are detected in the volume, a total of 3*M* key-frames will be returned by the algorithm. Note, that selecting the median and not the midpoint between the first and last B-scans, ensures that the B-scan included in the summary will be one that was identified by the deep learning classifier as being relevant for the visual summary.

**Algorithm 1:**
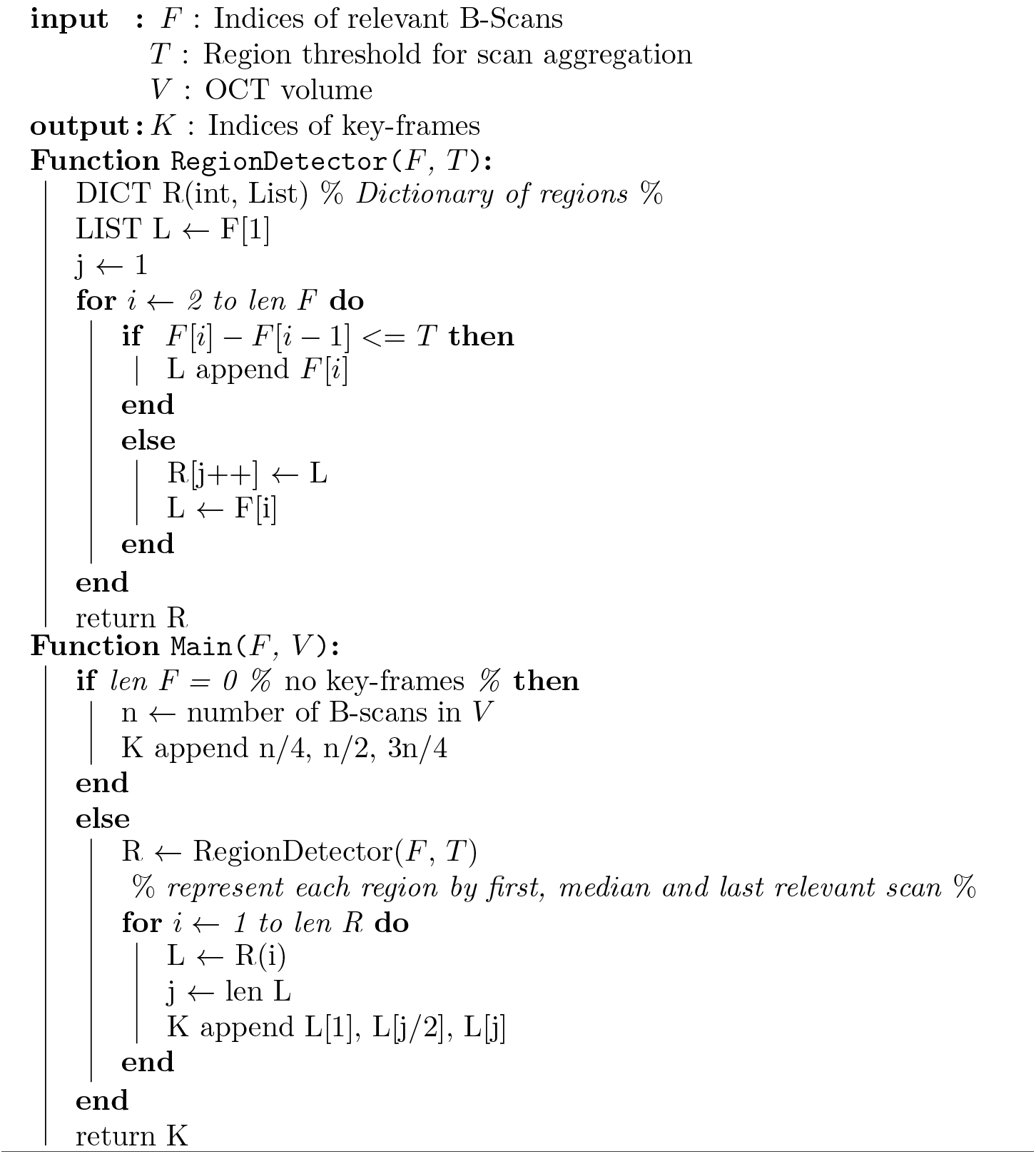
Summarisation Rules.

## 4 Results

The training of the transfer learning network required approximately 1000 epochs, while *de novo* training needed nearly 2000 epochs. This is to be expected as the transfer learning network is pre-trained. The initial loss was also substantially larger for the *de novo* network (Note the difference in *y*-axes scales in Fig. 5).

**Figure 5.**
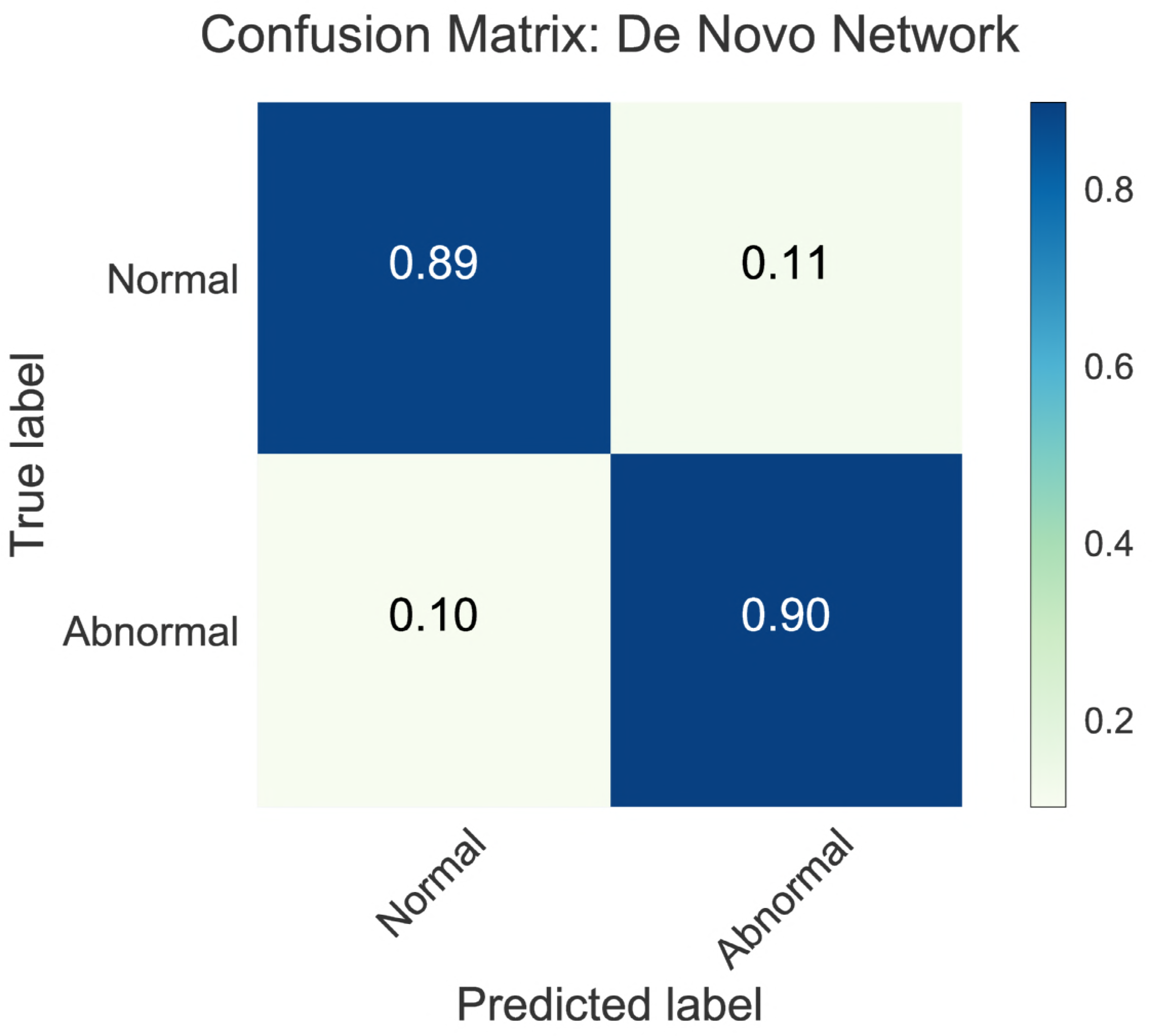
Loss and accuracy (AUC) monitored during (a) transfer learning and (b) *de novo* training.

The AUC, computed on the test set alone, was 0.97 and 0.96 for the transfer learning and the *de novo* networks, respectively (see Fig. 6(c)). The sensitivity (upper left quadrant of the confusion matrix) was found to be 0.91 for the transfer learning and 0.90 for the *de novo* network (see Fig. 6(a)-(b)). Similarly, the specificity was found to be 0.91 and 0.89 for the transfer learning and *de novo* networks, respectively. Fig. 7 shows instances of B-scans with mild ((a)-(c)) and severe ((d)-(f)) AMD-related pathologies that were correctly identified by the networks as being relevant to the visual summary. The CAM visualisation for mild and severe B-scans are shown in Fig. 7.

**Figure 6.**
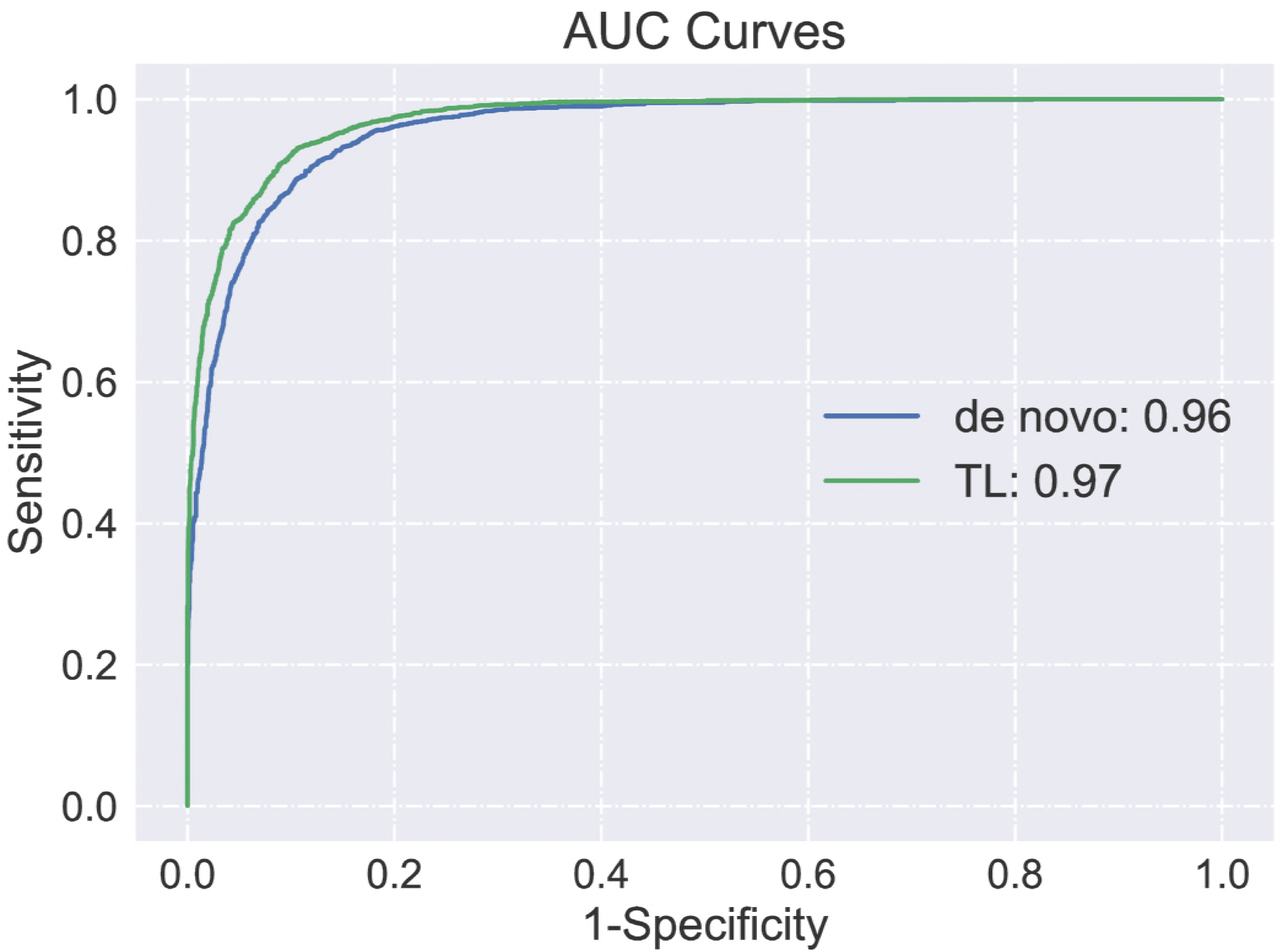
Confusion matrices for the (a) transfer learning (TL) and (b) the *de novo* network. (c) The AUC plot for the two networks.

**Figure 7.**
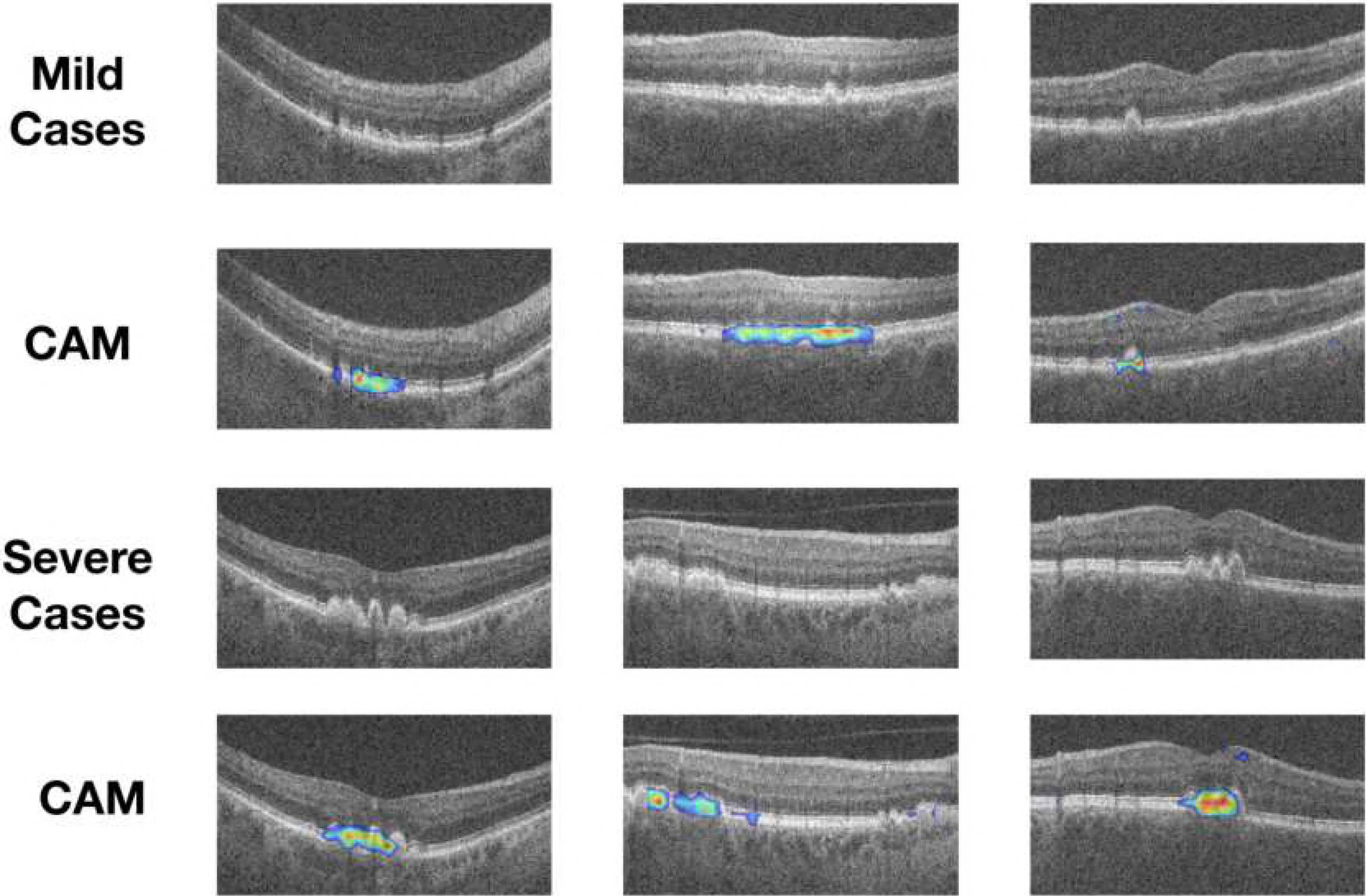
Examples of AMD B-scans with small/mild (top row) and significant pathologies (third row). CAMs from the *de novo* network for the same images are shown in the second and fourth rows, respectively.

The percentages of false positives arising from the control datasets was also computed, and found to be 6% and 7.5% for the transfer learning and *de novo* networks, respectively. Visualisations of these B-scans showed that poor quality scans from the healthy controls were sometimes misclassified. Errors in the retinal localisation as shown in Fig. 8(b) also led to misclassifications. The false negatives were analysed further in order to investigate why they were misclassified. Visually inspecting the results obtained from the transfer learning framework revealed that the false negatives in 73% of the cases contained small pathological conditions such as isolated drusen (see Fig. 8(c)). Similarly, for the *de novo* network, 91% of the false negatives showed “mild” visually discernible pathologies. However, there were also instances where geographic atrophy was not correctly identified as a pathology (see Fig. 8(d)). The dataset consists of horizontal as well as vertical scans [10], where normal B-scans close to the optic nerve head are visually very similar to geographic atrophy. Since the model did not incorporate this additional piece of information (horizontal or vertical scan), it is not surprising to see misclassifications of this particular pathology.

**Figure 8.**
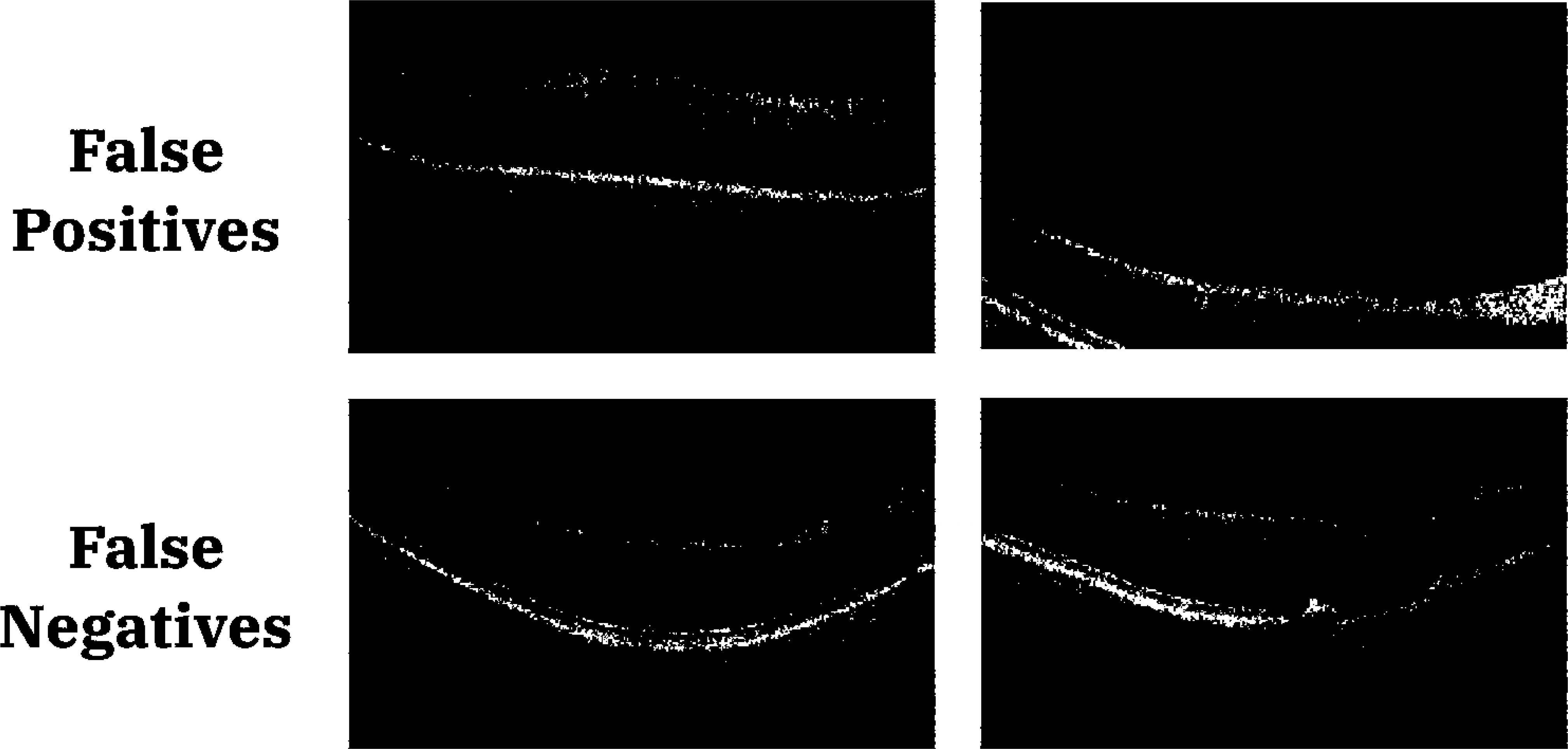
Examples of false positives from control subject scans (top row), and false positives in AMD scans (bottom row).

### 4.1 Summarisation Result

An example of a visual summary generated by the system is displayed in Fig. 9. This scan showed pigment epithelial detachment at the fovea, and only a single region was identified by the summarisation rules.

**Figure 9.**
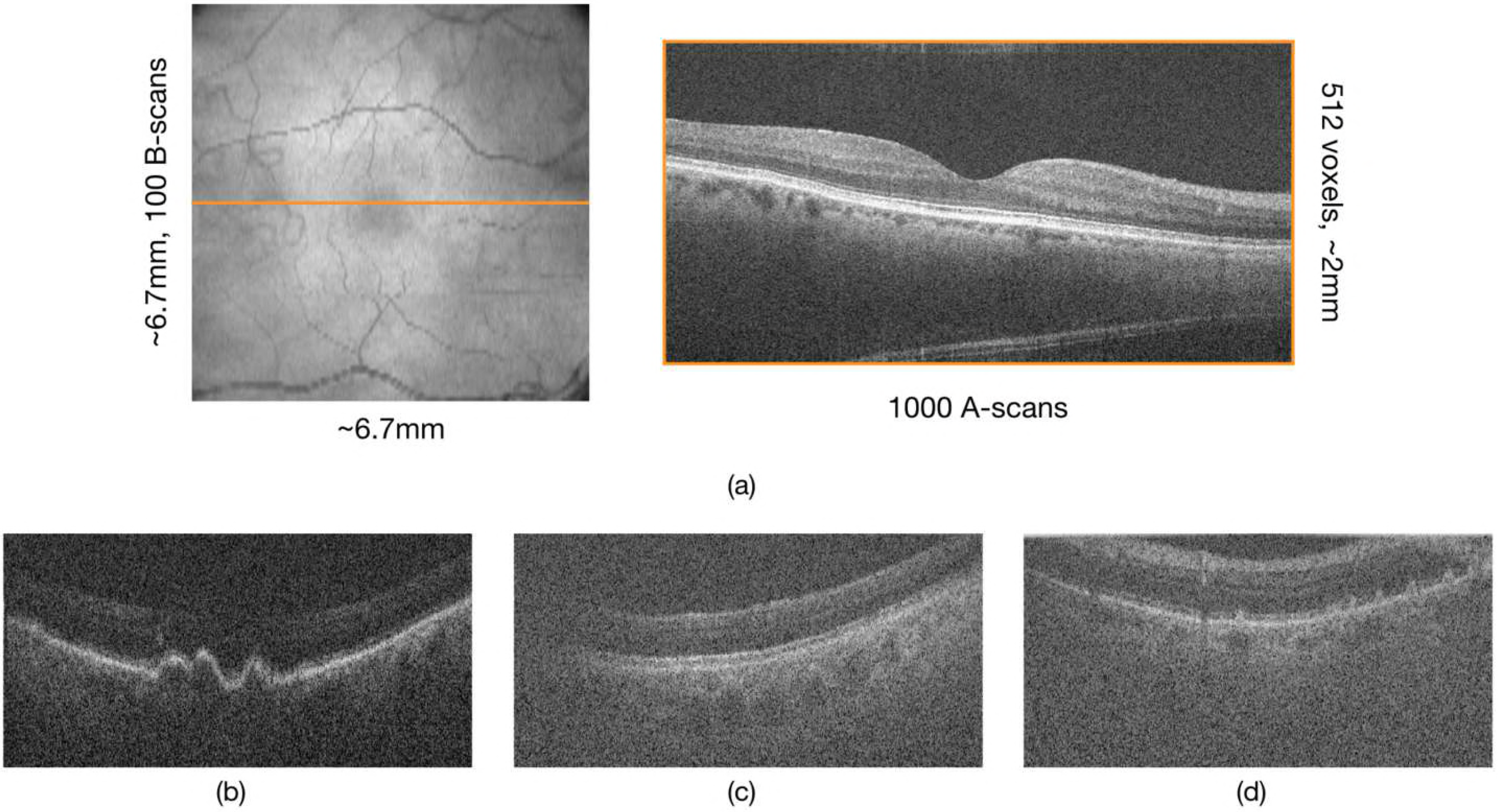
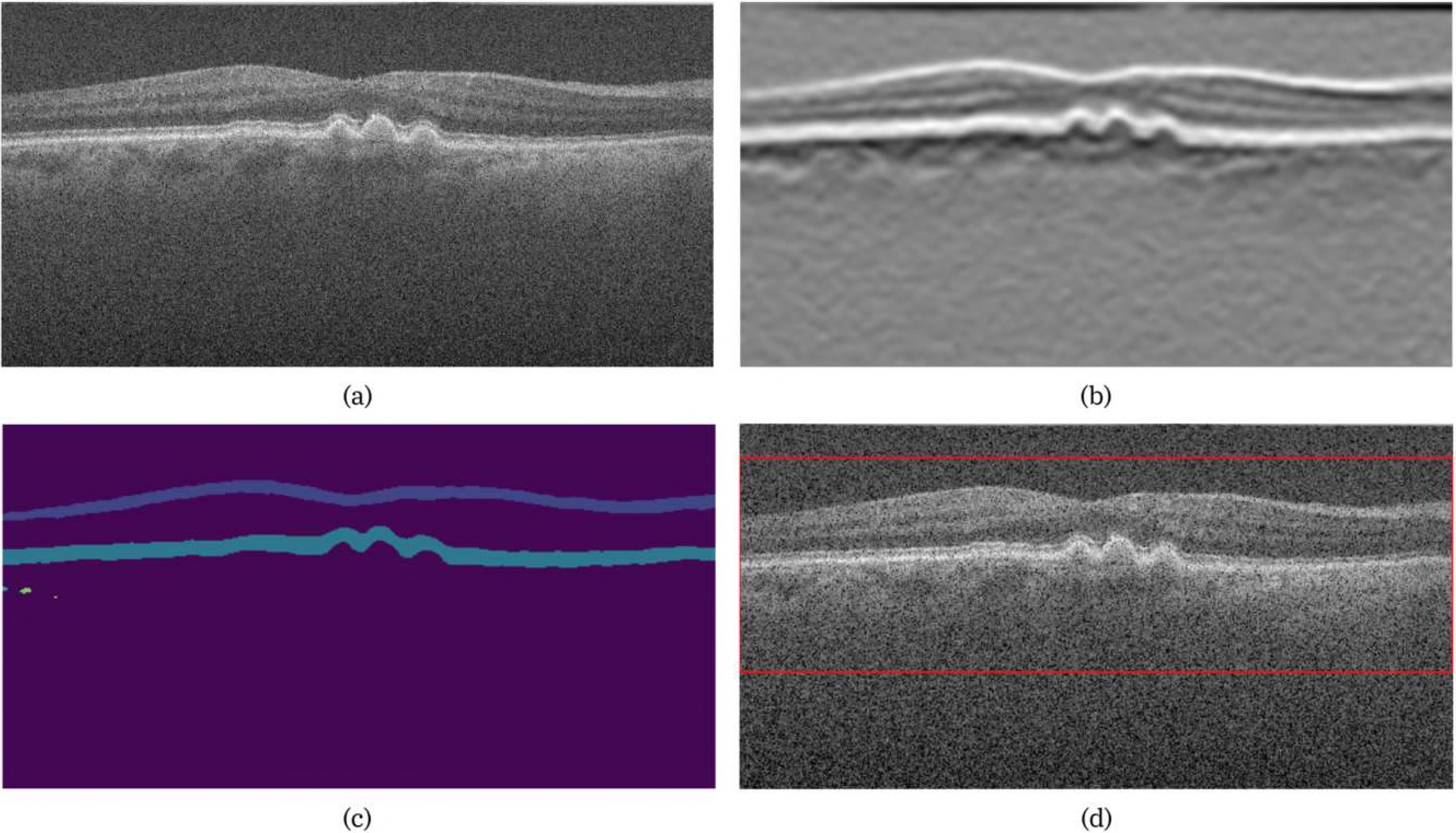
(a) The en-face projection image of the volume, with (b) all the key B-scans and (c) the final visual summary. The B-scans that correspond to the three locations are shown below, and colour coded to indicate their location in the volume.

A second example of an SDOCT volume with drusen and geographic atrophy is presented in Fig. 10. Here three separate regions were detected by the summarisation algorithm as indicated by the blue, red and green regions in Fig. 10(b). The final visual summary consisting of the B-scans that represent the three regions depicted in Fig. 10(c). The individual B-scans from the three regions are shown in the three rows Fig. 10(d) - (l), with the colours of the bounding boxes corresponding to the location indicated in Fig. 10(c).

**Figure 10.**
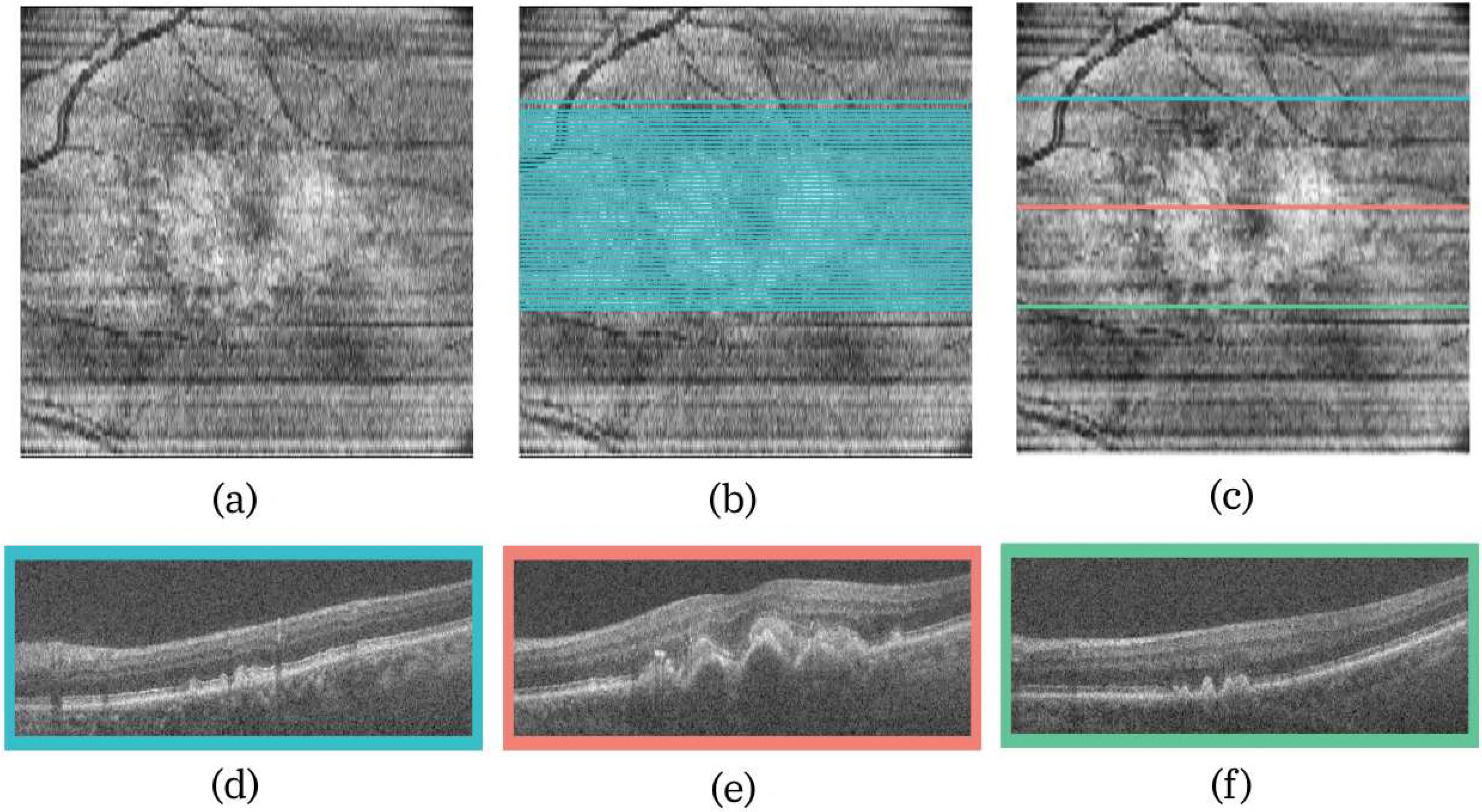
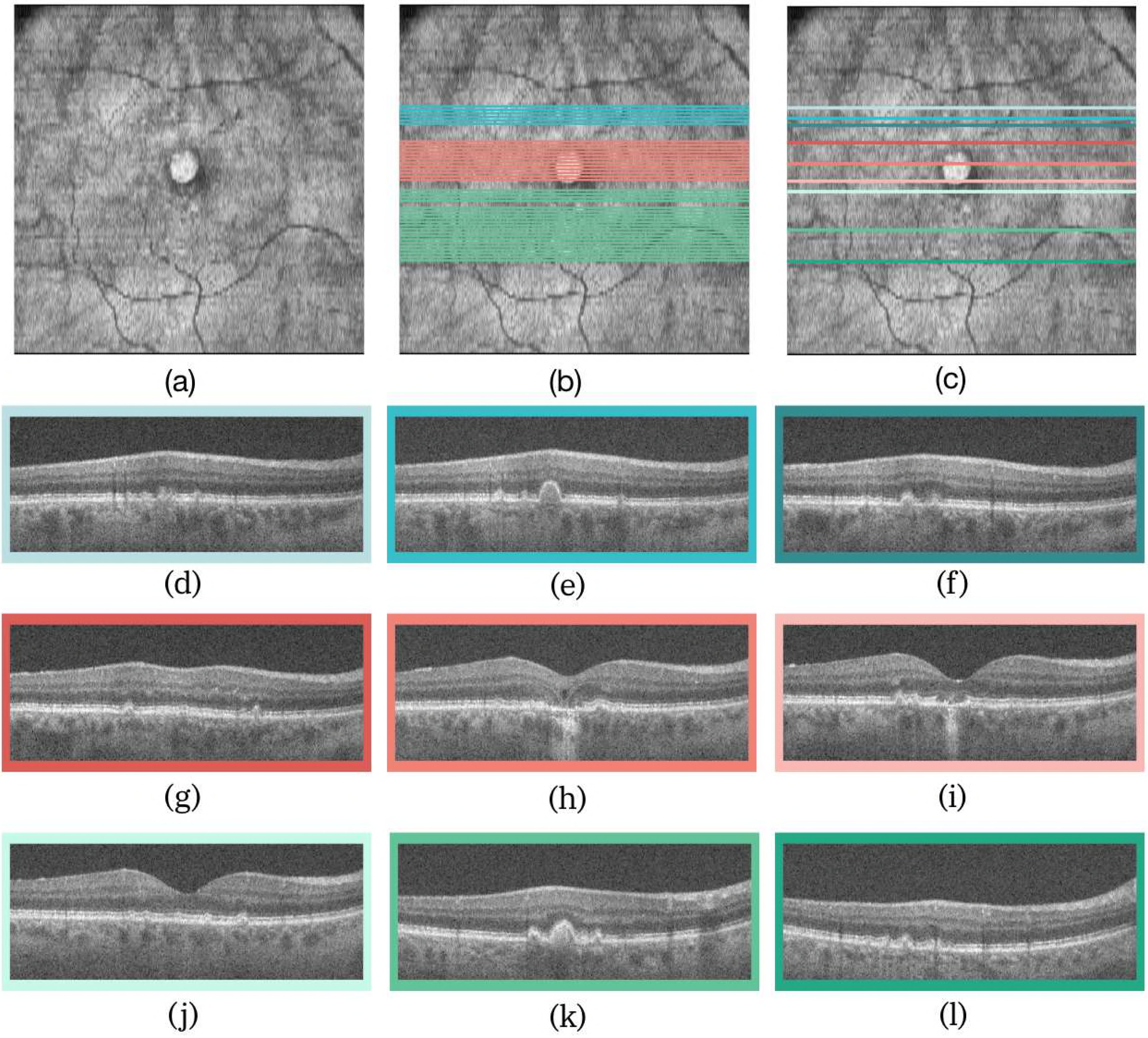
(a) The en-face projection image of the volume, with (b) all the key B-scans and (c) the final visual summary. The three B-scans corresponding (d) - (f) to the first region (blues), (g) - (i) the second region (reds), and (j) - (l) to the third region (greens).

## 5 Discussion & Conclusions

SDOCT finds extensive use in ophthalmology for the visualisation and quantification of structures in the retina. This high-resolution modality generates vast quantities of data (∼ 50MB per volume), making the visual inspection of these images time-consuming, tiring and therefore error prone. Summarisation of the SDOCT volumes has been limited to the extraction of structural measurements, such as retinal layer thicknesses or optic nerve head parameters such as cup-to-disc ratio. However, other conditions such as epiretinal membranes or intra-layer cysts (that do not affect retinal layer thickness) require the manual inspection of the B-scans in the OCT volume. Visual summaries that retrieve key B-scans and identify relevant regions of the scan can be a valuable addition to the existing diagnostic framework.

Previously proposed summarisation methods for videos or volumetric medical images rely on the detection of relevant features, similar to our approach. However, our method does not explicitly segment AMD-related pathologies (see [9]), but uses a deep learning network for the detection of B-scans that show structural abnormalities. Our method can be extended to any structural abnormality or even use a different definition of “relevance”. For instance, a similar system could be designed to extract B-scans where maximal temporal change is identified. This would allow to monitor a variety of conditions such as AMD (dry or wet) or even glaucoma, where changes at the optic cup are recorded over time. A system that analyses temporal SDOCT volumes, however, could not utilise transfer learning and would require a *de novo* network, designed and trained for the specific application. In our experiments, we found that the *de novo* training and network design can offer advantages over transfer learning, the most important being the relaxation of input restriction, e.g. three-channel input image of specific size vs a grayscale image of selectable size.

A separate experiment was conducted with the transfer learning network, where the input consisted of three adjacent B-scans instead of a replication of a single B-scan. Intuitively, one would expect that the use of adjacent B-scans would bolster the network’s ability to detect the key B-scans, but this network performed worse. The conclusion to be drawn is not that the adjacent B-scans have no additional useful information, but that the transfer learning network is ill-equipped to leverage this. The VGG-16 network was originally designed for three-channel colour images where each channel presented different colour characteristics of the same image. Here, the adjacent B-scans might show differing structures (healthy B-scans adjacent to one with small drusen), and the network was not able to efficiently leverage this additional information. A *de novo* network, designed and trained for this, might do better, but we did not pursue this.

The *de novo* network, being custom designed, also allowed for the incorporation of a the global average pooling layer (to generate the CAMs), as well as skip-connections (known to assist in training). The inclusion of the CAM brings a degree of “explainability” to the system, where this visualisation indicates the source of the final class label. This output could also be used as an input to the rules that generate the visual summaries, where the size of would CAMs impact the inclusion in the visual summary. The *de novo* network was 98% smaller than the VGG-16 network, but was found to be just as accurate and robust. This small size allowed for rapid training and required only two days (on a K80 NVIDIA GPU) instead of the six needed by the transfer learning network. Run-time is also affected by network size and the classification of an entire SDOCT volume only took four seconds for the *de novo* network, while the larger transfer learning network took 190 seconds (computed on a 2.5GHz Intel Core i7, 16GB RAM system).

In conclusion, the presented *de novo* network allows for the rapid and reliable detection of key B-scans in SDOCT volumes, which are used to generate visual summaries of volumes. In the future, we intend to extend the summarisation techniques to other disease models, as well as explore the use of small networks, designed specifically for the task at hand.

